# Neuronal Correlates of the Perceptual Invariance of Words and Other Sounds in the Supplementary Motor Area of Macaques

**DOI:** 10.1101/2020.12.22.424045

**Authors:** Jonathan Melchor, Isaac Morán, José Vergara, Tonatiuh Figueroa, Javier Perez-Orive, Luis Lemus

**Affiliations:** Department of Cognitive Neuroscience, Institute of Cell Physiology, Universidad Nacional Autónoma de México (UNAM). 04510. Mexico City, Mexico; Department of Neuroscience, Baylor College of Medicine, Houston, Texas; Luis Guillermo Ibarra Ibarra National Rehabilitation Institute. Mexico City, Mexico

**Keywords:** Perceptual constancy, macaques, acoustic recognition, frontal cortex, psychophysics

## Abstract

The supplementary motor area (SMA) of the brain is critical for integrating memory and sensory signals into perceptual decisions. For example, in macaques, SMA activity correlates with decisions based on the comparison of sounds.^1^ In humans, functional MRI shows SMA activation during the invariant recognition of words pronounced by different speakers.^2^ Nevertheless, the neuronal correlates of perceptual invariance are unknown. Here we show that the SMA of macaques associates novel sounds with behaviors triggered by similar learned categories when recognizing sounds such as words. Notably, the neuronal activity at single and population levels correlates with the monkeys’ behaviors (e.g. hits and false alarms). Our results demonstrate that invariant recognitions of complex sounds involve premotor computations in areas other than the temporal and parietal speech areas. Therefore, we propose that perceptual invariance depends on motor predictions and not only sensory representations. We anticipate that studies on speech will observe sensory-motor transformations of acoustic information into motor skills.

## INTRODUCTION

We typically learn our first words from the voice of our mother. Notably, after that, we are capable of recognizing the same words spoken by multiple speakers. Similarly, the ability to identify versions of learned sounds (vLS) has been studied in songbirds, ferrets, and mice.^3–5^Rhesus monkeys and chimpanzees acknowledge vocalizations for foods of different values and from various troop members.^6,7^ Although this ability is vital for communication, the neural basis for the invariant recognition of sounds is not well understood.

Current knowledge of how neuronal activity produces the semantic and invariant perception of vocal sounds is also scant.^4,8–10^ For instance, “voice-sensitive” and “vocalization-sensitive” cortical areas have been identified using neurophysiological and imaging studies in humans and nonhuman primates.^11–15^ Additionally, the invariant perception of ferrets generalizing vowel identities (/u/ and /ϵ/) throughout variations of fundamental frequencies, sound levels, and locations has been correlated with auditory cortex activity.^4^ In zebra finches, reports suggest invariant responses for different categories of conspecific vocalizations.^9^ A recent study in mice found that they discriminate the consonants (/g/ and /b/) and still recognize them when combined with different vowels or emitted by other speakers.^5^ Nevertheless, understanding the neuronal correlates of invariant perception of complex sounds such as words, requires active recognition during neurophysiological recordings.

Studies in primates have shown that auditory processing travels from the auditory cortex to the frontal lobe via the ventral auditory stream.^16,17^ In humans, the ventral stream groups syllables into words during speech recognition.^18,19^ The prefrontal cortex projects to premotor cortices, which orchestrate behaviors.^20^ For instance, the macaque ventral premotor cortex, homologous to Broca’s human speech production area, participates in acoustic discrimination.^21^ Also, as a premotor cortex, the SMA participates in voluntary movement control,^22–24^ working memory, and decision making.^25–28^ In humans, it is also involved in acoustic imagery.^20^ Recently, Vergara and collaborators showed that the SMA is crucial for auditory decisions^1^. Therefore, the SMA is a likely candidate area for linking sounds and the invariant perception thereof.^2^

Here we present extracellular recordings of SMA neurons in rhesus monkeys trained to discriminate sounds. During the task, the monkeys reported the appearance of a target sound (T) presented randomly after 0 to 2 non-target sounds (nT). We evaluated the neuronal responses to various vLS of T and nT to which the monkeys had had no previous exposure (e.g. a sound uttered by different emitters). We observed that the monkeys perceived vLS as the originally learned sounds (LS), and that the SMA’s neuronal responses were correlated with the animals’ invariant perceptions. Our results suggest that the SMA associates incoming unexperienced sounds with the closest known category.

## RESULTS

We trained two rhesus monkeys to categorize numerous sounds as T or nT during an acoustic recognition task (Fig. 1a; see Methods). The monkeys released a lever for a T in the first, second, or third position of a sequence (Fig. 1b). Trials with one, two, or three stimuli had an equal probability of appearing in a session (p = 1/3). To assess the monkeys’ capacity to identify each sound, we computed the probability of hits p(release | T) and correct rejections CR = p (no-release | nT) (Fig. 1b, inset). Overall, each monkey performed above chance for more than 20 sounds,^29^ [Hits median: monkey1 = 0.97, monkey2 = 0.96; CR median: monkey1 = 0.98, monkey2 = 0.96; one-sample Wilcoxon signed-rank test higher than 0.75: Z (monkey1_Hits) = 10.41, Z (monkey1_CR) = 8.51, Z (monkey2_Hits) = 9.63, Z (monkey2_CR) = 7.87; all p-values < 0.001]. Overall, performance for all T and nT categories ranked above 85% for both monkeys. The monkeys learned 36 sounds (T = 11, nT = 25). The sounds consisted of seven artificial sounds (T = 2, nT = 5), six monkey vocalizations (T = 2, nT = 4), sixteen human words (T =6, nT = 10), and seven other animal vocalizations (T = 1, nT = 6; see reference 29 for further behavioral details).

**Figure 1.**
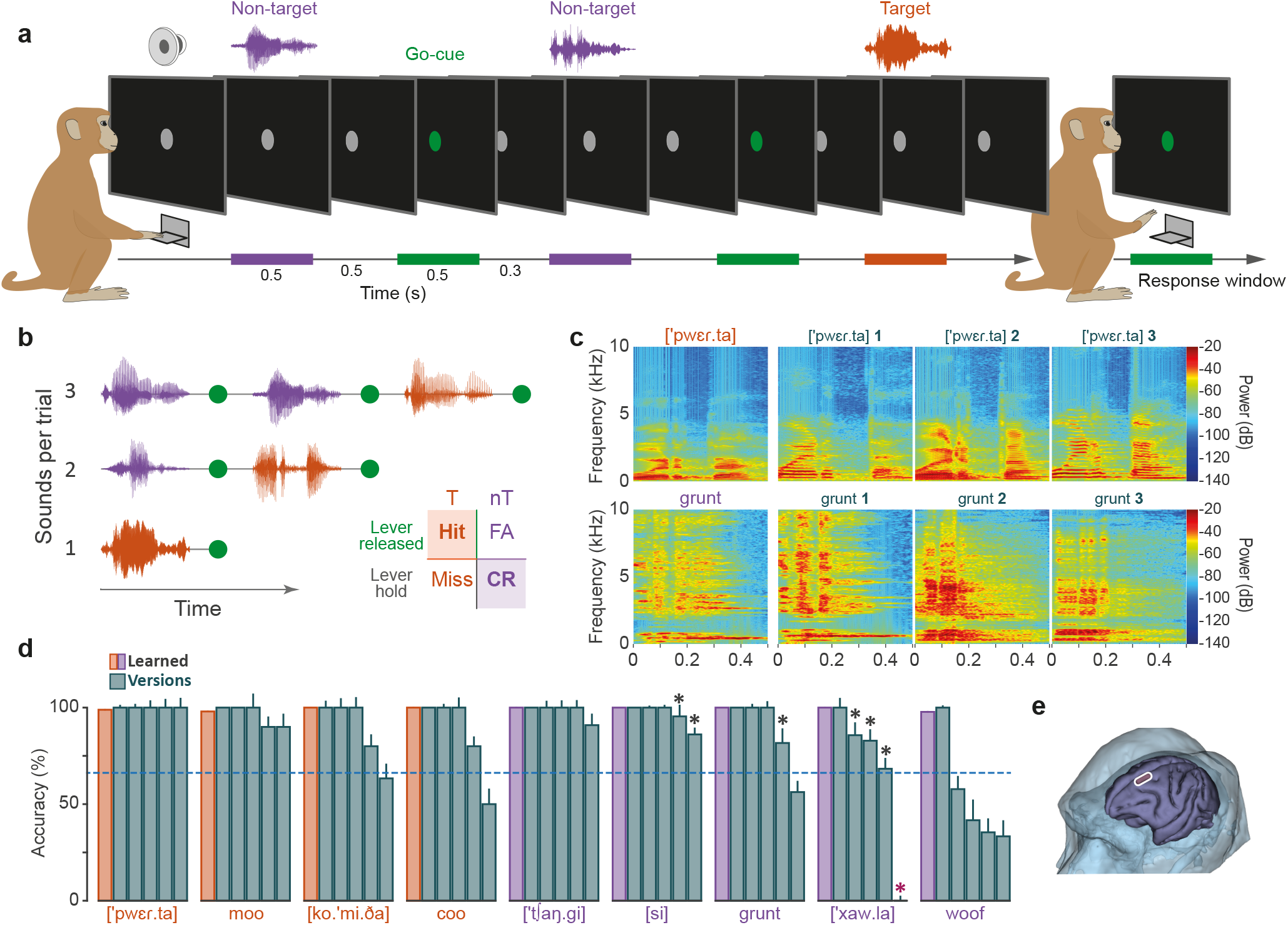
Auditory recognition task and behavioral performance. **a)**Events in a trial. First, a visual cue (grey circle) appears at the center of the screen, indicating that the monkey should press and hold down the lever. After a variable period (0.5 to 1 s), a playback of 1-3 sounds commenced. Each sound continued with a 0.5 s delay and a 0.5 s green cue that replaced the grey circle. The monkey received a reward for releasing the lever within 0.7 s after the beginning of the green cue. Color code: orange (T), purple (nT), green (go-cue). **b)**Types of trials. The T could appear in the first, second, or third position during a trial. Inset: Behavior as a function of the lever release in response to T and nT. FA, false alarms; CR, correct rejections. **c)**Spectrograms of T and nT sounds and some acoustic versions (upper and lower rows, respectively). **d)**Mean hit rates for learned T and CR for learned nT (orange and purple bars, respectively) and vLS (grey bars). The central horizontal line demarcates acoustic recognition by chance at 50%. Black asterisks: performance different from LS. Red asterisk: an exceptional sound that was recognized as a T. **e)**Structural MRI of monkey 1; SMA indicated.

To test for invariant recognition of sounds, we presented the monkeys with several vLS. Figure 1c shows the T [‘pwεr.ta] and the nT ‘grunt’spectrograms, together with three vLS. To reduce the possibility of learning the vLS, we presented each intercalated with LS no more than fifteen times per session. Ultimately, the monkeys recognized significantly above chance 84.4% of vLS (n = 45; Wilcoxon signed-rank test; Benjamini-Hochberg FDR corrected q-value = 0.01). This result means that the animals succeeded in categorizing vLS as either T or nT. However, to verify whether the monkeys perceived vLS as an invariant of LS, we sought performance differences between each of the nine LS and their corresponding five vLS (Fig. 1d). Overall, the monkeys recognized 71% vLS of LS (pairwise multiple Wilcoxon rank-sum comparison tests, Benjamini-Hochberg FDR correction, q value = 0.01). It is noteworthy that the ‘woof category was highly biased towards false alarms during nT vLS since the monkeys only recognized 1 out of 5 vLS for the LS ‘woof.’ Overall, with the exceptions of the vLS ‘woof and the vLS [‘xaw.la], the monkeys invariantly perceived most vLS as LS.

Once the monkeys were performing the task consistently, we recorded 65 SMA neurons (Fig. 1e) in order to determine whether the neural responses showed invariance to vLS. Figure 2a shows the raster plots of a neuron’s responses to LS and vLS (left and right panels, respectively). Here, the peristimulus time histograms (PSTHs) in Fig. 2b describe similar response patterns of increased firing rates in all T regardless of whether they were LS or vLS. Interestingly, the neuron fired after the vLS ‘woof up to the activity level of the T (dashed line). Then, at the beginning of the visual cue, the firing rate fell to nT levels. Notably, neither the monkeys nor the neurons correctly classified the ‘woof sound as nT.

**Figure 2.**
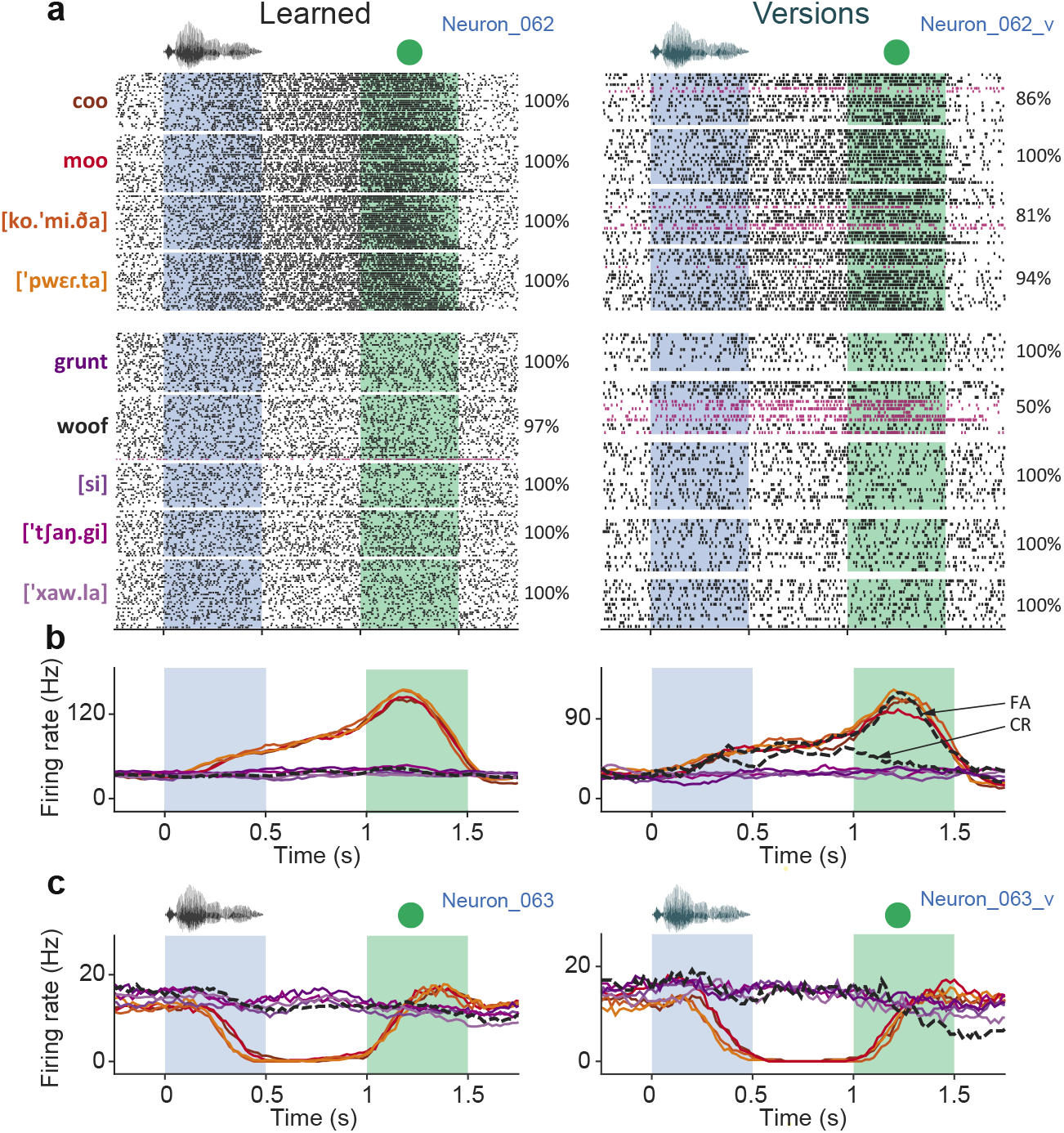
Responses of SMA neurons during the recognition of learned and version sounds. **a)**Raster plot of a neuron’s responses during trials of LS and vLS (left and right panels, respectively). Each line is a trial aligned to the beginning of sounds. Each dot is an action potential. The sounds were presented randomly during the experiment but shown here in blocks of T and nT sounds (upper and lower groups, respectively. **b)**PSTHs of the neuron in **a**. CR, correct rejections, and FA, false alarms of vLS ‘woof.’**c)**PSTHs of a second neuron recorded in the same session.

During that experimental session, we also recorded the activity of other neurons, e.g. the neuron in Figure 2c. As opposed to the neuron described above, this neuron inhibited its responses to all T to almost zero spikes/second. However, it was unaffected by nTs, including vLS of ‘woof,’ which means that in this case the excitatory and not the inhibitory activity was biased during the false alarms.

So that we might evaluate the extent to which the neuronal population also coded T and nT as invariant, i.e. regardless of their particular identities, we performed an *F*-statistic analysis. Figure 3a shows the number of neurons coding for sounds or categorical decisions during LS or vLS. Fifty-three out of the 65 recorded neurons (81.5%) showed categorical responses to LS and 49 to vLS (75.4%). However, 2 neurons (3%) coded for a particular T or nT LS and 6 (9%) for nT vLS. Figure 3b presents the periods in which the neuronal population codes for T and nT in LS and vLS. Interestingly, the f-statistic values for LS categorical responses were higher than vLS, thus showing a more robust difference between T and nT rather than an effect of the number of neurons which was almost the same for LS and vLS conditions.

**Figure 3.**
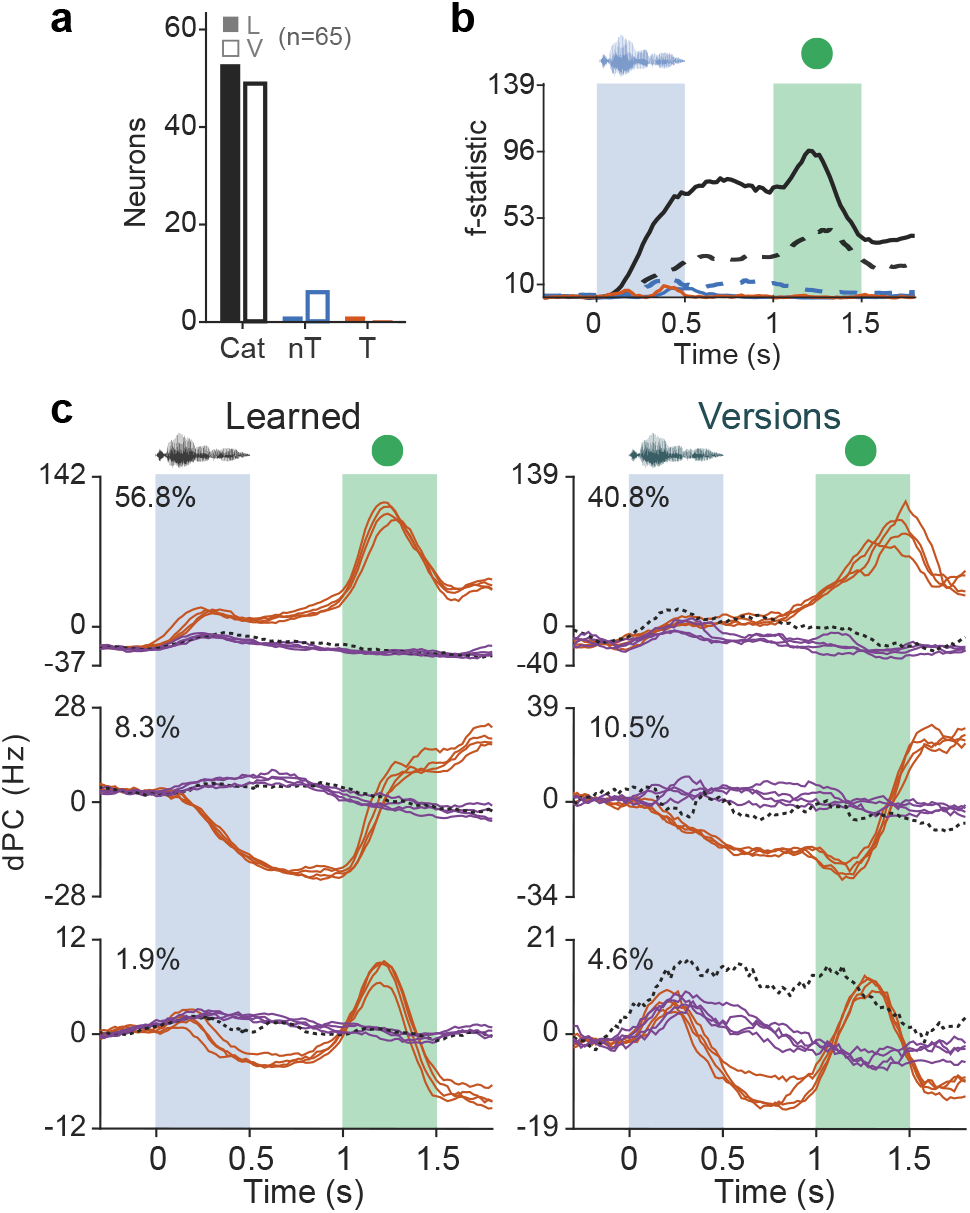
Categorical responses of the SMA population. **a)**Neurons with categorical responses to T or nT groups or particular identities during LS and vLS computed by *F*-tests. **b)**SMA mean *F*-statistic values as a function of time. Black lines: categorical responses. Orange: T identities. Blue: nT identities. Continuous lines: LS. Dashed lines: vLS. **c)**Each row: first three demixed principal components of the SMA neuronal population during LS and vLS. Dashed line: ‘woof sound. Orange colors: T. Purple colors: nT.

We also performed dPCA (see methods) in order to verify whether the neuronal population can code for acoustic identities within categorical representations. The population activity observed with dPCA showed similar patterns for LS and vLS (Figure 3c). Here, the three principal components reflect categorical coding of T. However, this analysis did not show tuning to particular T or nT sounds. One exception occurred at the third component of vLS where it was observed that the ‘ woof’ stands out from the nT distribution. As described in Figure 2b, the neuronal population seems to explain the monkey’s false alarms. These results suggest that invariant perception in individual and population neuronal activity does not depend on recognition of the timbre or modulations of particular emitters. Instead, invariant perception relies on a more general representation of each category.

In this regard, to test whether the neuronal responses were correlated with the monkeys’ behavior, we calculated Spearman correlations between behavior and the firing rates of LS and vLS in 200 ms bins calculated every 20 ms (Fig. 4a). Figure 4b shows the significant Spearman correlations throughout the task’s periods and each neuron. The mean population’s rho values varied similarly over time (i.e. Pearson’s R = 0.90, p < 0.001). This implies that the neuronal population relates the task’s events with the monkey’s behavior for both LS and vLS. Moreover, the neurons that engaged across the events showed similar LS and vLS dynamics (Figure 4c; Pearson’s R = 0.86, p < 0.001). Overall, these results suggest that individual and population neuronal activity correlates with invariant perception in rhesus monkeys.

**Figure 4.**
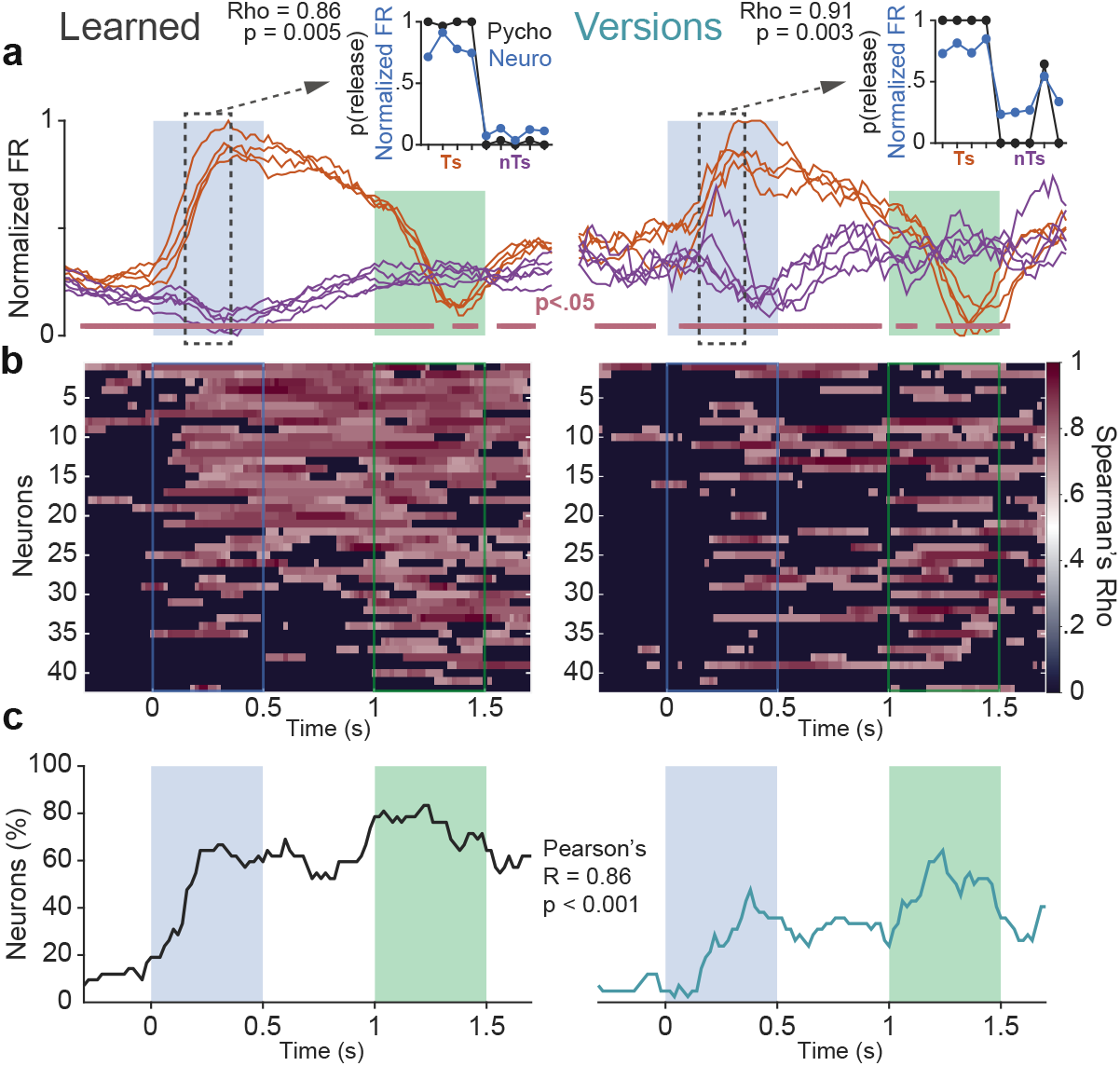
Correlation between the monkeys’ performance and SMA neuronal responses. **a)**Responses of 1 neuron to LS and vLS. Insets: a time window indicated by the dashed box, where the probability of the monkey releasing the lever after each T and nT (black line) is similar to neuronal responses to LS and vLS (blue line). Rho: the Spearman’s correlation and p significance for that temporal window. **b)**Significant Spearman’s rho along with the task for each neuron during LS and vLS. **c)**Times where the neuronal population recorded during LS and vLS were correlated with the monkey’s performance.

## DISCUSSION

Our results demonstrate that rhesus monkeys correctly categorized sounds to which they had had no previous exposure as different variants of learned sounds. This result is significant because in nonhuman primates, vocal production is highly stereotyped and probably genetically determined.^30–34^ Numerous studies report limited acoustic recognition in nonhuman primates^35–37^; however, emerging evidence suggests long-term plasticity in vocal learning in adult monkeys exposed to vocalizations during development.^38,39^ For example, in a match-to-sample task, macaques were better at conspecific calls than other vocalizations or sounds.^40^ Although we used similar sounds, we did not find performance biases towards conspecific vocalizations. One possible explanation is that with the experiments of Ng et al., discrimination of monkey vocalizations was better because those sounds had existed in long-term memory since birth. In contrast, working memory alone was probably more demanding and required different brain circuits. In our task, there was no working memory involved. After the monkeys learned numerous sounds that may or may not be ethologically relevant, they practiced for many days until they consistently achieved performance above 90%. Therefore, we expect different processing of short and long-term memories in primates.^41–43^ Future studies may give insight into those differences.

Although there are no reports of perceptual invariance of sounds in non-human primates, our results show that the behavior and SMA responses to LS and vLS were invariant. In other words, the SMA activity supports the constant perception of sounds. This result is similar to experiments in primary auditory areas demonstrating perceptual invariance during the recognition of vowels.^4^ However, our work does not show that the SMA represents particular sounds but rather that it directs actions based on acoustic recognition. Notably, those sounds are recognized when they relate to the motor actions triggered by known sounds. In other words, the SMA relates sounds to behaviors.^1^ Thus, it is reasonable to suggest that the SMA is able to sort various types of sensory information regardless of whether or not it is stored in long term circuits.

In our task, different acoustic categories lead to the release or the holding down of the lever. Therefore, each group of heterogeneous sounds produces 1 of 2 learned motor outputs. Arguably, many sounds become synonyms of “release the lever” and many others of “hold the lever.” In agreement with this fact, SMA neuronal responses were correlated with two orthogonal motor plans emerging from the recognition of T or nT sounds. Noticeably, responses were invariant to T or nT groups and not to particular acoustic categories within each group. Experiments in visual areas such as the inferotemporal cortex have demonstrated that single neurons represent visual categories regardless of sensory modulations or visual perspective.^44,45^ Therefore, instances in the hierarchical processing of sensory information create invariant representations of images. However, in those experiments, the neurons were not tested during active recognition. Thus, perceptual invariance was not tested. In contrast, experiments with monkeys that actively categorized visual^46–48^ and auditory stimuli^49–51^ showed that neuronal activity in the prefrontal cortex was correlated with the animals’ performance. These results suggest that areas in the frontal lobe are responsible for perceptual experiences while sensory areas create invariant sensory signals. Further experiments will disentangle whether perceptual invariance is created at higher cortical areas fed by invariant sensory representations.

## CONCLUSIONS

In rhesus monkeys, the invariant perception of sounds is more than recognition of physical identities of sounds, e.g. particular timbres and modulations. It also relies on the motor representations linked to each category. In this sense, our results suggest that the SMA is a critical instance for the auditory invariant perception of sounds such as words, because it associates un-experienced acoustic stimuli with the categorical responses triggered by known sounds.

## METHODS

### Ethics statement

All experimental protocols were performed in compliance with the Official Mexican Standard Recommendations for the Care and Use of Laboratory Animals (NOM-062-ZOO-1999) and approved by the Internal Committee for the Use and Care of Laboratory Animals at the Institute of Cell Physiology, UNAM (CICUAL; LLS80-16).

### Animals and experimental setup

Two adult rhesus macaques *(Macaca mulatta;* one 13 kg, ten year old male, and one 6 kg, ten year old female) participated in this study. The animals inhabited an enriched facility that allowed physical interactions with other monkeys. The monkeys were restricted to water only the night before experimental sessions. However, after the sessions, they received water *ad libitum.* Monkeys performed ~1000 trials during three-hour sessions, at one session per day, four to five days per week. Experiments took place in a soundproof booth in which a macaque sat on a primate chair, 60 cm away from a 21” LCD color monitor (1920 x 1080 resolution, 60 Hz refreshing rate). A Yamaha MSP5 speaker (50 Hz-40 kHz frequency range) set 15 cm above and behind the monitor delivered acoustic stimuli at ~65 dB SPL measured at the monkey’s ear level. Additionally, a Logitech^®^ Z120 speaker was positioned directly below the Yamaha speaker in order to render white background noise at ~55 dB SPL. Finally, a metal spring-lever at the monkeys’ waist level captured their responses.

### Behavioral task

We trained two rhesus macaques in an acoustic recognition task that consisted of categorizing sounds presented in trials of 1 to 3 sounds as target (T) or non-target (nT). At each trial, a 3°-aperture gray circle appeared at the monitor’s center, after which the monkey pressed the lever. After a variable delay of 0.5 to 1 s, a 0.5 s sound was heard, followed by a 0.5 s delay and a 0.5 s period when the gray circle turned green. If the sound consisted of a T, the macaque released the lever within a 0.8 s response window commencing with the green cue. However, if the sound was an nT, the monkey kept the lever down and waited for the next sound. We presented the T at the 1st, 2nd, or the 3rd position with equal probability, i.e. p(T | position) = 1/3. Releases before a T created false alarms, leading to different new trials. We required that the monkeys perform above an 80% hit rate before electrophysiological recordings. The task’s programming was in LabVIEW 2014 (64-bit SP1, National Instruments®).

### Acoustic stimuli

The sounds were recorded in our laboratory or downloaded from free online libraries (https://freesound.org/). They consisted of words, monkey vocalizations, other animal vocalizations, and artificial sounds (n = 37, Supplementary Table 1). The sample rate was 44.1 kHz; cutoff frequencies, 100 Hz to 20 kHz, compressed or extended to 0.5 s, and equalized to the same root-mean-square (RMS) amplitude value (Adobe Audition® version 6.0). We selected 4 T and 5 nT from the pool of learned sounds in order to perform statistical repetitions (ten times each sound in a set). The phonetic nomenclature of Spanish words proceeded from the automatic phonetic transcriptionist created by Xavier López Morrás (http://aucel.com/pln/transbase.html). To assess the monkeys’ perceptual invariance to unheard sounds, we presented them with 5 versions of each learned sound.

### Neuronal recordings

We performed extracellular recordings of single SMA neurons in monkey 2. We positioned a 20 mm diameter recording chamber above the stereotaxic coordinates of the SMA (Paxinos G 2009) compared to monkey 2 MRI (IA = 27 mm, left hemisphere: lateral 6 mm). We used an array of 5 independently movable microelectrodes (1-3 MΩ; Thomas Recording®) inserted at different locations in each session. We sampled extracellular membrane potentials at 40 kHz and performed offline sorting using Plexon sorter software (Plexon®).

### Analysis

To assess the monkeys’ capacity to identify each sound, we computed the probability of hits [p(release | T), T] and Correct Rejections [CR = p(no-release | nT), nT], we performed a one-sample Wilcoxon signed-rank test. To verify the difference between the LS and vLS, we calculated the number of vLS that were different from LS (pairwise multiple Wilcoxon ranksum comparison tests, Benjamini-Hochberg FDR correction, q value = 0.01).

To evaluate what aspects of the task were coded by SMA neurons, e.g. T and nT categories or acoustic identities, we calculated *F*-statistics in a one-way ANOVA of the T vs. nT and each sound. We calculated the mean firing rate in 200-ms windows in steps of 20 ms. In order to avoid biases in the *F*-statistic due to the number of trials, we only included the same number in all classes. We repeated the analysis 1000 times to create an *F*-statistic distribution. We compared the actual distribution against an *F*-statistic random distribution obtained by shuffling the stimulus labels 1000 times to determine the significant *F*-statistic bins. A neuron was significant if confidence intervals (95%) did not overlap in at least one time-bin.

We performed demixed principal component analysis (dPCA)^52^ in order to observe the effects of the task components and of acoustic identities on the population responses. During the supervised stage, dPCA decomposes the neural activity for each variable in covariance matrices of different marginalizations. Then, for each marginalization matrix, unsupervised analysis is similar to a PCA. We marginalized the population activity for T, nT, and time, from the beginning of the sounds to the monkeys’ responses, in 200 ms windows every 20 ms. dPCA differentiates the matrices from the decoder (D) and the encoder (F) for each parameter by minimizing the loss function:

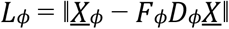

Where *φ* denotes the marginalization for each parameter and *X*, the mean-centered population matrix. Each component represents an amount of firing rate variance. The axes obtained by the decoder and the encoder allow data reduction in a few features, capturing the most significant variance of each parameter.

To determine statistically decoded bins, we organized the data into training and testing sets. The testing sets consisted of random inclusions of 1 trial from each category (T and nT). The training set was the mean of the remaining tests of each class. We performed dPCA to the training set in order to obtain the decoding axes. Then, we projected the testing set on these axes and classified them according to the closest category mean (T or nT). We repeated this procedure 1000 times and measured the proportion of correct classification. We shuffled 100 times for significance.

To test whether the neuronal responses were correlated with behavior, we computed Spearman correlations between the monkeys’ performance and neuronal firing rates of LS and vLS, in 200 ms windows, every 20 ms. Finally, we calculated a Pearson correlation between the resulting mean Spearman’s rho of LS and vLS in order to assess for similarities.

## ACKNOWLEDGMENTS

We are grateful for the financial support provided by CONACYT **CB-256767**, and *Programa de Apoyo a Proyectos de Investigatión e Innovatión Tecnológica* [Support Program for Research Projects and Technological Innovation *(PAPIIT)***IN207919**. Special thanks to Gerardo Coello and Ana Escalante of the computing department of the IFC and to Patrick Weill for proofreading the manuscript. Jonathan Melchor Hernández is a doctoral student in the Programa de Doctorado en Ciencias Biomédicas [doctoral program in biomedical sciences], at Universidad Nacional Autónoma de México (UNAM) and he was supported by CONACYT 229866. The data in this work are part of his doctoral dissertation.

## AUTHOR CONTRIBUTIONS

JM, IM, TF, and LL performed experiments, JM, IM, JV, and JP analyzed data, LL designed the paradigm, TF programmed the task, while JM and LL wrote the paper.

## CONFLICT OF INTEREST STATEMENT

The authors declare no competing financial interests.

## Notes

### Competing Interest Statement

The authors have declared no competing interest.

